# Associations between striatal neurochemical changes and resting-state functional connectivity following transcranial temporal interference stimulation

**DOI:** 10.64898/2026.09.02.749008

**Authors:** Shota Kimura, Hiroyuki Matsuta, Kana Matsumura, Fumiaki Iwane, Nobuhiro Hata, Yoshiki Asayama, Hisato Sugata

## Abstract

**Background:** Non-invasive modulation of deep brain structures remains a major challenge in human neuroscience and neurorehabilitation. Transcranial temporal interference stimulation (tTIS) has emerged as a promising approach for engaging subcortical regions, but its effects on striatal neurochemical markers and functional connectivity remain unclear.

**Objective:** To investigate whether 20-Hz tTIS designed to target the right striatum modulates GABA+ and glutamate–glutamine complex (Glx) levels and whether neurochemical changes are associated with changes in resting-state functional connectivity.

**Methods:** Thirty-four healthy right-handed participants were randomly assigned to the tTIS group (20-Hz beat frequency) or sham group (0-Hz frequency difference). Participants underwent proton magnetic resonance spectroscopy (^1^H-MRS) and resting-state fMRI before and after 30 min of stimulation. One participant in the tTIS group was excluded from the ^1^H-MRS analyses because of data corruption.

**Results:** The direct between-group difference in Glx change did not reach statistical significance, although the tTIS group showed a numerically greater reduction than the sham group (p = 0.066). Exploratory within-group analyses showed a significant decrease in striatal Glx in the tTIS group (p < 0.001), whereas the decrease in the sham group was not significant. No significant GABA+ changes were observed within or between groups. The association between striatal Glx change and functional connectivity change differed between groups in the sensorimotor cortex, right parahippocampal gyrus, and bilateral fusiform gyri.

**Conclusion:** These exploratory findings suggest that 20-Hz tTIS may be associated with striatal Glx modulation and group-dependent coupling between neurochemical and cortico-subcortical network changes.

## 1. Introduction

Non-invasive brain stimulation techniques are widely used to investigate and modulate human brain function, as well as to enhance motor skill learning ^1^. Their widespread use stems from their non-invasive nature, which allows them to be applied relatively safely and easily in both healthy and clinical populations ^2–5^. Although conventional non-invasive brain stimulation techniques such as TMS and tDCS primarily modulate cortical regions, their ability to focally target deep brain structures is limited ^6, 7^. In contrast, deep brain stimulation (DBS) has demonstrated efficacy in treating neurological disorders such as Parkinson’s disease and epilepsy ^8, 9^. However, DBS requires surgical implantation and is therefore invasive, substantially limiting its use in healthy participants and its applicability to basic neuroscience research ^10^.

These limitations have prompted the development of alternative non-invasive neuromodulation techniques. One emerging approach is transcranial temporal interference stimulation (tTIS). tTIS is thought to preferentially modulate deep brain structures while minimizing the direct recruitment of overlying cortical neurons ^11^. The mechanism underlying tTIS is based on the interference of two high-frequency electric fields with slightly different frequencies, which produce a low-frequency envelope of amplitude modulation within the region of field overlap ^11,12^. Recent human studies have shown that tTIS can modulate activity in deep brain structures such as the striatum and hippocampus ^13–15^. Because the basal ganglia play a central role in motor control ^16^, neuromodulation of basal ganglia circuits may have therapeutic potential for neurological disorders such as Parkinson’s disease. Recent pilot studies have begun to explore the clinical potential of tTIS, and have reported improvements in motor symptoms in patients with Parkinson’s disease following stimulation of basal ganglia structures ^17^. Beyond its potential motor applications, tTIS is also being investigated for its effects on neuropsychiatric conditions. For example, stimulation targeting the nucleus accumbens has been reported to improve depressive symptoms in patients with bipolar disorder, suggesting that tTIS may modulate functionally relevant deep brain circuits beyond the motor domain ^18^.

Neural oscillations are organized into distinct frequency bands, including alpha, theta, and beta rhythms, each of which is associated with different cognitive and sensorimotor functions ^19–21^. Among these rhythms, beta oscillations are considered a prominent feature of sensorimotor network activity. In particular, beta oscillations in sensorimotor regions are thought to support the maintenance of the current sensorimotor state, consistent with the idea that beta-band activity reflects the status quo of sensorimotor processing ^22^. Throughout the sensorimotor system, beta oscillations around 20 Hz are prominently observed across multiple components involved in movement control and somatosensory processing, including muscles, dorsal root ganglia, basal ganglia, and cerebral cortex ^23^. Beta-band oscillations are present throughout the basal ganglia and are modulated during the processing of sensory cues and motor-related signals ^24^. Importantly, beta oscillations do not arise from a single structure but instead reflect coordinated activity across the cortico-basal ganglia network. Thus, modulating beta oscillations around 20 Hz represents a physiologically grounded strategy for modulating cortico-basal ganglia network dynamics.

Because beta-band activity is closely linked to cortico-basal ganglia network function, understanding this circuit requires consideration of the neurochemical architecture of this circuit. Within cortico-basal ganglia circuitry, cortical information is processed through two principal striatal pathways: the direct and indirect pathways, which have traditionally been associated with movement facilitation and suppression, respectively ^25, 26^. However, both pathways can be concurrently activated, and their balanced activity contributes to action selection and movement generation ^27, 28^. Beta-band oscillations may contribute to this coordination by stabilizing the current motor set rather than promoting the initiation of new movements ^22^. At the neurochemical level, corticostriatal projections are predominantly glutamatergic, whereas striatal projection neurons of both the direct and indirect pathways are GABAergic ^25^. The interplay between glutamatergic corticostriatal input and GABAergic striatal output is therefore central to the regulation of basal ganglia function ^25, 26^.

Although previous studies have demonstrated that tTIS can modulate deep brain activity and associated functional networks ^13–15^, it remains unclear whether tTIS alters neurochemical markers within deep brain structures. In particular, little is known about how tTIS influences neurochemical indices related to excitatory and inhibitory neurotransmission in the striatum. Therefore, we hypothesized that tTIS targeting the striatum would alter striatal GABA and the glutamate–glutamine complex (Glx) levels, and alter striatal resting-state functional connectivity. To test this hypothesis, we investigated whether 20-Hz tTIS targeting the striatum alters striatal GABA and Glx levels using proton magnetic resonance spectroscopy (^1^H-MRS), and whether stimulation-induced changes in these metabolites are associated with alterations in resting-state functional connectivity measured by functional magnetic resonance imaging.

## 2. Methods

### 2.1. Ethics statement

This study protocol was reviewed and approved by the Ethics Review Board of the Oita University Faculty of Welfare and Health Science (approval number: F250012) and conducted in accordance with the Declaration of Helsinki. All participants were fully informed about the purpose and procedures of the study and provided informed consent before participation.

### 2.2. Participants

34 healthy young adults participated in this study (mean age, 20.8 ± 1.7, range, 19–29 years). The inclusion criteria were (i) right-handedness and (ii) age between 18 and 29 years. Exclusion criteria included the presence of a cardiac pacemaker or metallic implant, claustrophobia, and a history of neurological or psychiatric disorders. All participants were confirmed to be right-handed using the Edinburgh Handedness Inventory (EHI) ^29^. The mean EHI score of the tTIS group was 87.9±16.9, and the mean EHI score of the sham group was 89.8 ± 12.9 (Table 1). Following enrollment, participants were randomly assigned to the tTIS or the sham group.

**Table 1.** Characteristics of the participants in each group.

|  | **tTIS group** | **sham group** | ***p* value** |
| --- | --- | --- | --- |
| Gender (male/female) | 10 / 7 | 11 / 6 | 0.72 |
| Age (mean ± SD) | 20.6±0.89 | 20.7±2.2 | 0.86 |
| EHI score (mean ± SD) | 87.9±16.9 | 89.8±12.9 | 0.72 |

### 2.3. Transcranial Temporal Interference Stimulation (tTIS)

Transcranial temporal interference stimulation (tTIS) was applied non-invasively using a tTIS device (Soterix Medical, Inc., USA). In both groups, the electrode pairs were positioned at F3-F4 and TP7-TP8 to target the right striatum, in accordance with a previous study ^15^. Participants in the tTIS group received sinusoidal electrical currents oscillating between −2 and +2 mA in each stimulation channel, at carrier frequencies of 2,000 and 2,020 Hz. These stimulation currents generated a 20-Hz amplitude-modulated envelope in the basal ganglia. In contrast, participants in the sham group received sinusoidal currents of the same amplitude, with a carrier frequency of 2,000 Hz in both stimulation channels. Thus, the frequency difference was 0 Hz, producing no low-frequency amplitude-modulated envelope (Fig. 1A). The stimulation period consisted of a 10-second ramp-up, 30 min of continuous stimulation, and a 10-second ramp-down, resulting in a total stimulation duration of 30 min 20 s (Fig. 1B).

**Figure 1.**
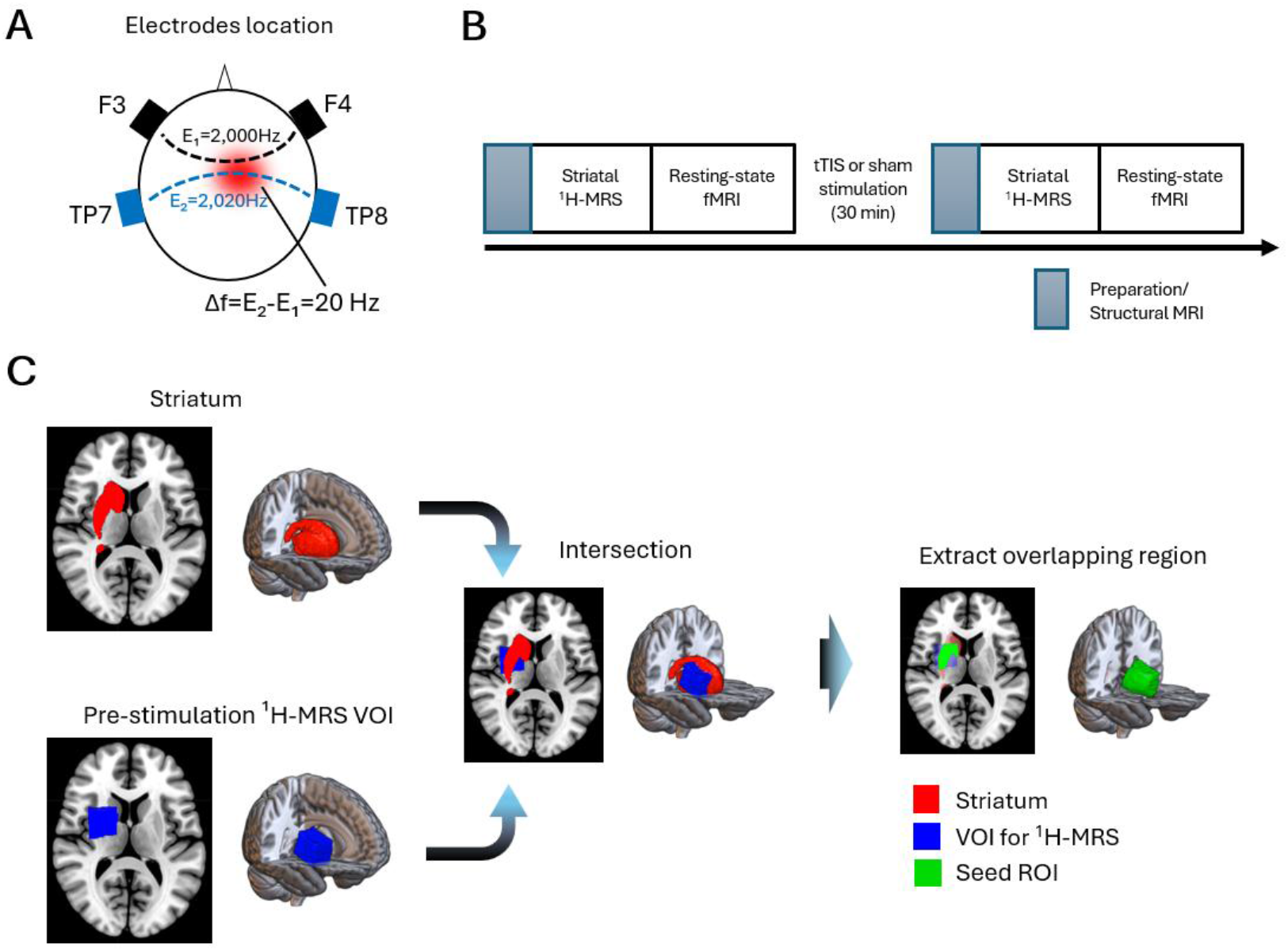
Experimental design and definition of the striatal seed ROI. (A) Electrode locations and stimulation parameters. Electrodes were placed at F3, F4, TP7, and TP8, forming the F3–F4 and TP7–TP8 electrode pairs. In the tTIS condition, sinusoidal currents at 2,000 and 2,020 Hz were delivered through the two electrode pairs, respectively, producing a 20-Hz temporal interference envelope intended to target the right striatum. In the sham condition, the same electrode montage was used, but both currents were delivered at 2,000 Hz, resulting in a frequency difference of 0 Hz and no low-frequency interference envelope. (B) Experimental procedure. Participants underwent preparation and structural MRI, striatal ^1^H-MRS, and resting-state fMRI before and after the intervention. The intervention consisted of 30 min of either tTIS or sham stimulation. (C) Definition of the subject-specific striatal seed ROI. Each participant’s pre-stimulation ^1^H-MRS VOI was spatially aligned with the anatomical right striatal mask. The intersection between the pre-stimulation ^1^H-MRS VOI and the right striatal mask was extracted and used as the subject-specific striatal seed ROI for seed-based functional connectivity analysis. The same seed ROI was used for both the pre- and post-stimulation resting-state fMRI sessions. Red indicates the right striatal mask, blue indicates the pre-stimulation ^1^H-MRS VOI, and green indicates the extracted seed ROI.

### 2.4. Proton magnetic resonance spectroscopy (^1^H-MRS) and functional magnetic resonance imaging (fMRI) acquisition

Before ^1^H-MRS acquisition, high-resolution structural images were acquired on a 3-T MRI scanner equipped with a 32-channel head coil (MAGNETOM Skyra Fit, Siemens Healthineers, Germany) using a T1-weighted three-dimensional MPRAGE sequence. T1-weighted images were acquired in the axial plane and reconstructed in the coronal and sagittal planes to guide the placement of a striatal volume of interest (VOI). The imaging parameters were as follows: repetition time (TR) = 2,500 ms, echo time (TE) = 2.18 ms, inversion time (TI) = 1,000 ms, flip angle = 8°, field of view (FOV) = 240 × 256 mm, matrix size = 300 × 320, and voxel size = 0.8 × 0.8 × 0.8 mm³.

Proton magnetic resonance spectroscopy (^1^H-MRS) data were then acquired from a striatal VOI measuring 30 × 25 × 25 mm³. MEGA-PRESS-edited spectra for the quantification of GABA+ and the glutamate–glutamine complex (Glx) ^30–34^ were acquired using the following parameters: repetition time (TR), 3 s; echo time (TE), 68 ms; editing pulses applied at 3.0 ppm during ON acquisitions and at 6.4 ppm during OFF acquisitions; 2,048 data points per acquisition; spectral bandwidth, 2 kHz; and 64 ON and 64 OFF acquisitions obtained in an interleaved manner. In addition, eight dummy scans were acquired for signal normalization. A total of 136 scans were acquired from each participant, resulting in an acquisition time of 408 s. Water suppression was achieved using the VAPOR sequence. Before spectral acquisition, manual shimming was performed over the VOI to achieve a water-peak full width at half maximum (FWHM) of ≤20 Hz.

After ^1^H-MRS acquisition, resting-state fMRI data comprising 375 volumes were acquired using a T2*-weighted single-shot gradient-echo echo-planar imaging (EPI) sequence. Each volume comprised 60 axial slices with a slice thickness of 2.4 mm and no slice gap, providing coverage of most of the cerebrum and cerebellum. EPI scans were acquired with the following parameters: repetition time (TR) = 800 ms, echo time (TE) = 34.4 ms, flip angle = 52°, field of view (FOV) = 206 mm, voxel size = 2.395 × 2.395 × 2.4 mm³, and matrix size = 86 × 86. The total acquisition time was 5 min. During resting-state fMRI acquisition, participants were instructed to remain still and fixate on a fixation cross presented in front of them.

All T1-weighted MRI, ^1^H-MRS, and fMRI data were acquired before and after the tTIS or sham stimulation session by H.M., a neurosurgeon with extensive experience in ^1^H-MRS acquisition (Fig. 1B).

### 2.5. ^1^H-MRS analysis

The ^1^H-MRS data were imported into the spectral analysis software LCModel (version 6.3-1N) ^35, 36^. After frequency and phase drift correction across the 64 ON and 64 OFF acquisitions, the data were Gaussian-filtered at 2 Hz and Fourier transformed before spectral quantification. Using an unsuppressed water-reference scan, eddy-current correction and water scaling were performed. GABA+, a composite signal comprising GABA and co-edited macromolecular contributions, and Glx, the combined signal of glutamate and glutamine, were quantified from the MEGA-PRESS-edited spectra using LCModel. Because the absolute concentrations of GABA+ and Glx are difficult to determine, GABA+/NAA and Glx/NAA ratios were calculated using the NAA signal obtained from the same acquisition and are hereafter referred to as the “GABA+ level” and “Glx level,” respectively. The MRS VOI masks were then coregistered to each participant’s T1-weighted structural image, and the structural images were segmented into gray matter (GM), white matter (WM), and cerebrospinal fluid (CSF) using Gannet (version 3.1.5) ^37, 38^. Because metabolite levels differ among GM, WM, and CSF, tissue correction was applied based on the tissue composition of each VOI.

### 2.6. VOI analysis

To confirm that the ^1^H-MRS VOIs were consistently positioned in the same brain regions across participants and between the pre- and post-stimulation scans in both groups, the VOIs were spatially normalized to Montreal Neurological Institute (MNI) space using SPM25 (https://www.fil.ion.ucl.ac.uk/spm/). The spatial overlap between the pre- and post-stimulation VOIs was visualized using MRIcroGL (https://www.nitrc.org/projects/mricrogl). For each participant, the pre-to-post VOI overlap rate was calculated as the number of voxels shared by the pre- and post-stimulation VOIs divided by the total number of voxels in the pre-stimulation VOI, and multiplied by 100 to express the value as a percentage. The resulting overlap rates were compared between the tTIS and sham groups using an independent-samples Student’s t-test to assess whether the consistency of VOI placement differed between groups.

### 2.7. Resting-state functional connectivity analysis

Resting-state functional connectivity analyses were performed using the CONN toolbox ^39^ running in MATLAB (MathWorks, USA). Seed-based connectivity analyses were conducted using the right striatum as the seed region. In the primary analysis, a subject-specific seed ROI was defined as the intersection between each participant’s pre-stimulation ^1^H-MRS VOI and the right striatal mask (Fig. 1C). The same seed ROI was used for both the pre- and post-stimulation resting-state fMRI sessions within each participant to avoid confounding longitudinal changes in connectivity with differences in seed location. Right striatal GABA+ and Glx levels were quantified using ^1^H-MRS before and after stimulation. The pre-to-post change in each ^1^H-MRS-derived neurochemical measure was evaluated separately. Any measure showing a significant pre-to-post change was subsequently entered as a neurochemical regressor of interest in the functional connectivity analysis. Pre-to-post changes in seed-based functional connectivity were estimated for each participant.

In the second-level analysis, each selected ^1^H-MRS-derived neurochemical change measure was entered as a regressor of interest in the CONN general linear model. Group-specific regressors were created for the tTIS and sham groups, allowing the association between neurochemical change and functional connectivity change to be estimated separately for each group. The difference in this association between groups was then evaluated by comparing the regression slopes for the tTIS and sham groups. Given the exploratory nature of the neurochemical–connectivity association analysis, statistical maps were thresholded at an uncorrected voxel-level threshold of p < 0.0005, combined with a cluster-extent threshold of k > 30 voxels. This exploratory thresholding strategy was adopted based on previous fMRI and resting-state functional connectivity studies that reported findings using uncorrected voxel-level thresholds together with minimum cluster-size or cluster-extent criteria of approximately 30 voxels ^40–42^. Clusters meeting these exploratory thresholds were interpreted as regions in which the relationship between the pre-to-post neurochemical change and the pre-to-post change in right striatal seed-based functional connectivity differed between the tTIS and sham groups.

## 3. Results

### 3.1 Side effects of tTIS and sham stimulation

No serious adverse events occurred during or after either tTIS or sham stimulation. Among the 34 participants, 17 reported slight pain (mean VAS score ± SD, 11 ± 14.5), 9 reported itching (9.1 ± 15.9), and 15 reported mild unpleasantness (11 ± 17.6).

### 3.2 Changes in striatal GABA+ and Glx levels

Due to data corruption during acquisition, one participant in the tTIS group was excluded from the subsequent analyses. Before evaluating the neurochemical changes, we compared the pre-to-post VOI overlap rates between the tTIS and sham groups (Fig. 2A). The comparison showed no significant between-group difference, indicating that the consistency of VOI placement was comparable between the groups (*t*(31) = 0.098, *p* = 0.923, *Cohen’s d* = 0.0341) (Fig. 2B). In all remaining participants, GABA+ and Glx signals were successfully quantified in the right striatum at both the pre- and post-stimulation time points (Fig. 2C). A direct between-group comparison of the percentage change from pre- to post-stimulation revealed no significant group difference in GABA+ change (*t*(31) = 1.375, *p* = 0.179, *Cohen’s d* = 0.479, two-tailed). The group difference in Glx change did not reach statistical significance, although the tTIS group showed a numerically greater reduction than the sham group (*t*(31) = −1.909, *p* = 0.066, *Cohen’s d* = −0.665, two-tailed). Exploratory one-sample t-tests against zero showed no significant change in GABA+ in either the tTIS group (*t(*16) = 1.578, *p* = 0.134, *Cohen’s d* = 0.383) or the sham group (*t*(15) = −0.237, *p* = 0.816, *Cohen’s d* = −0.059). In contrast, Glx decreased significantly relative to baseline in the tTIS group (*t*(16) = −8.113, *p* < 0.001, *Cohen’s d* = −1.968), whereas the reduction in the sham group did not reach statistical significance (*t*(15) = −1.846, *p* = 0.085, *Cohen’s d* = −0.462) (Fig. 3). These within-group findings were exploratory and should be interpreted in light of the nonsignificant direct between-group comparison of Glx changes.

**Figure 2.**
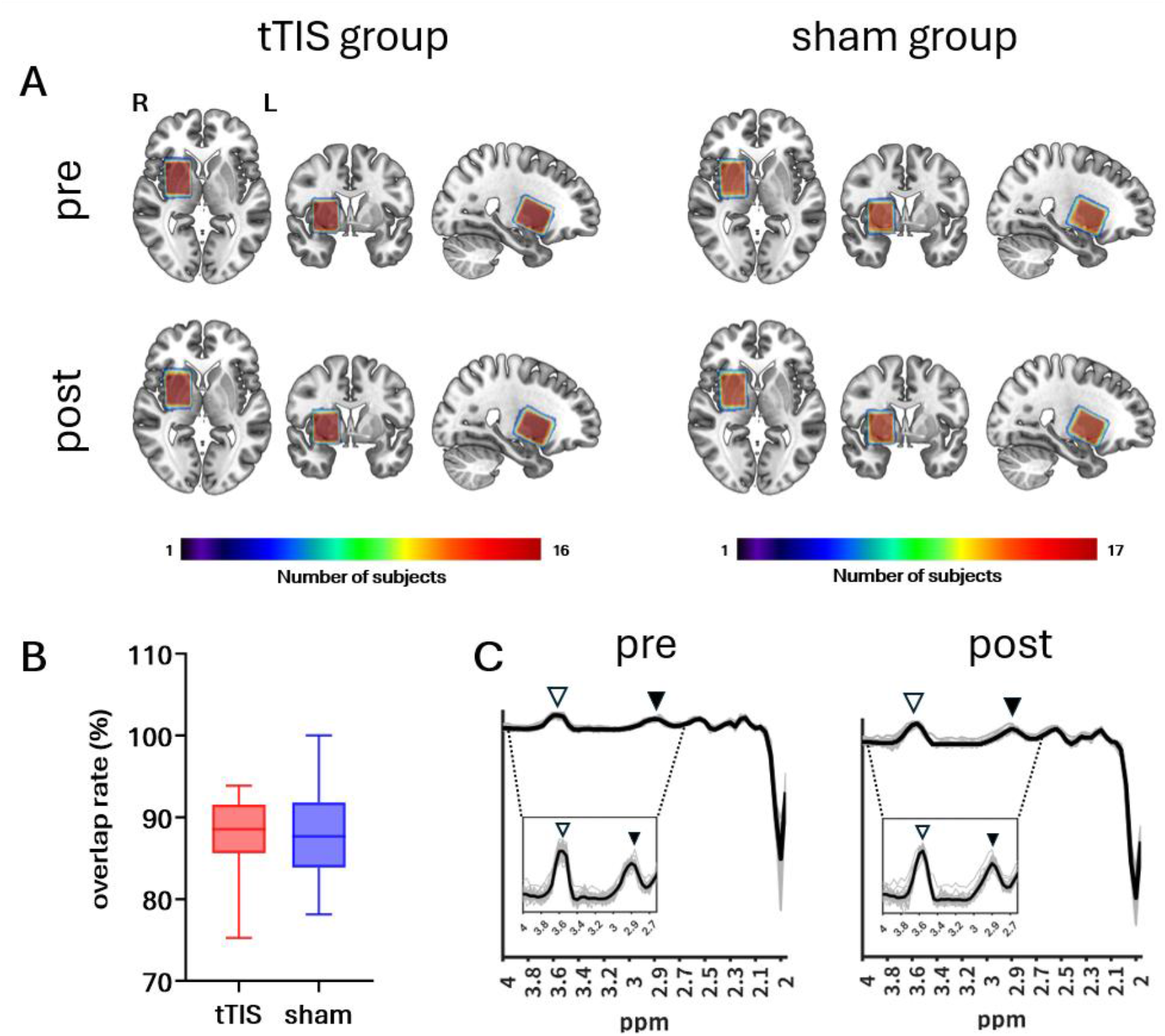
Spatial distribution and pre-to-post overlap of ^1^H-MRS VOIs and acquired spectra. (A) Spatial distribution of the ^1^H-MRS volumes of interest (VOIs). VOIs from all participants included in the ^1^H-MRS analysis were overlaid separately for the tTIS and sham groups at the pre- and post-stimulation time points. The color scale indicates the number of participants whose VOIs included each voxel within each group. (B) Pre-to-post overlap rate of the ^1^H-MRS VOIs. Box plots show the percentage of voxels in the pre-stimulation VOI that were also included in the post-stimulation VOI for each participant in the tTIS and sham groups. Red and blue indicate the tTIS and sham groups, respectively. (C) Individual and mean ^1^H-MRS spectra. Spectra from all participants included in the analysis were overlaid across both groups separately for the pre- and post-stimulation time points. Gray lines indicate individual participant spectra, and black lines indicate the mean spectrum across participants. Insets show expanded spectral regions highlighting the Glx signal at approximately 3.6 ppm (white arrowheads) and the GABA+ signal at approximately 2.9 ppm (black arrowheads).

**Figure 3.**
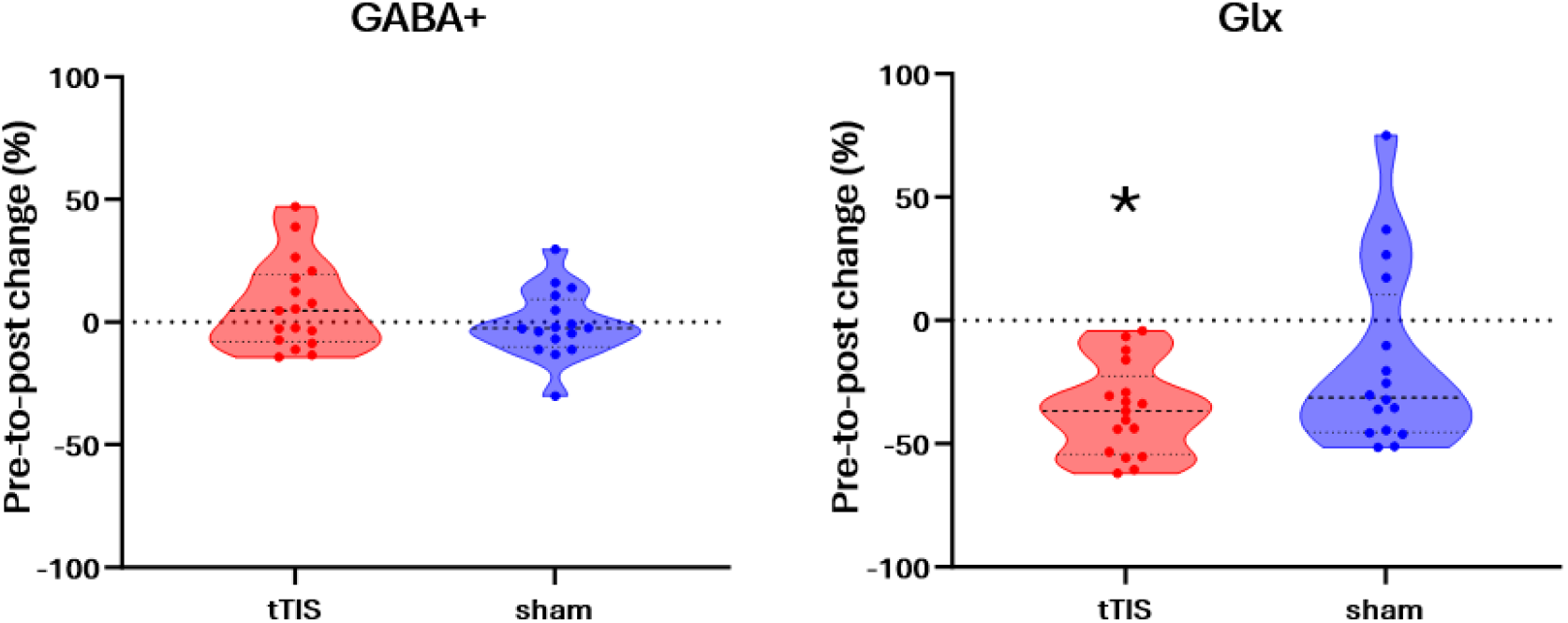
^1^H-MRS-derived changes in right striatal GABA+ and Glx levels. Percentage changes in right striatal GABA+ and Glx levels from pre- to post-stimulation are shown for the tTIS and sham groups. The percentage change in Glx was significantly below zero in the tTIS group, indicating a significant pre-to-post reduction. The asterisk indicates statistical significance based on a one-sample t-test against zero (*p* < 0.001). No significant between-group differences were observed in the percentage changes in either GABA+ (*p* = 0.179) or Glx (*p* = 0.066). The dotted horizontal line indicates zero change from baseline.

### 3.3 Association between striatal Glx changes and functional connectivity changes

Given the exploratory within-group decrease in Glx observed in the tTIS group, we next examined whether individual differences in right striatal Glx change were associated with pre-to-post changes in resting-state functional connectivity from the right striatal seed. The association between the percentage change in Glx and the change in connectivity strength differed between the tTIS and sham groups in several cortical clusters. Clusters meeting the exploratory thresholds were identified in sensorimotor-related cortical regions, including the right precentral and postcentral gyri, as well as in the right parahippocampal gyrus and bilateral fusiform gyri (Fig. 4A, Table 2). Across these clusters, the regression slopes were negative in the tTIS group and positive in the sham group. Thus, in the tTIS group, greater reductions in right striatal Glx tended to correspond to greater increases in connectivity strength, whereas the sham group showed the opposite pattern (Fig. 4B). These exploratory findings indicate that the relationship between right striatal Glx change and right striatal seed-based functional connectivity change differed between the tTIS and sham groups.

**Figure 4.**
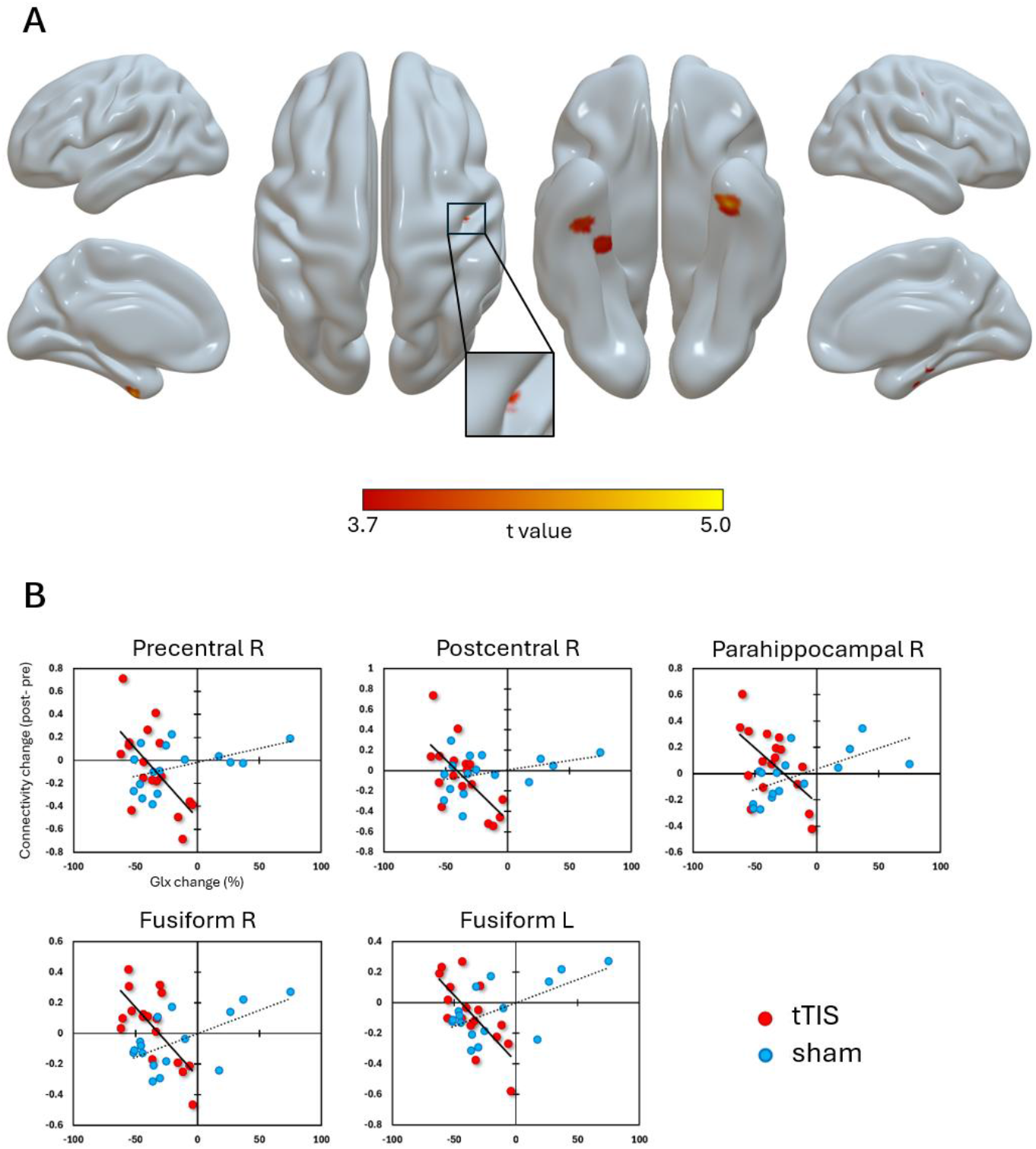
Group differences in the association between right striatal Glx changes and functional connectivity changes. (A) Clusters showing group differences in the association between the percentage change in right striatal Glx and the pre-to-post change in seed-based functional connectivity from the right striatal seed. Clusters meeting the exploratory statistical thresholds were identified in sensorimotor-related cortical regions, including the right precentral and postcentral gyri, as well as in the right parahippocampal gyrus and bilateral fusiform gyri. Statistical maps are displayed on cortical surface renderings, with the color scale indicating t values for the between-group difference in regression slopes. (B) Scatter plots showing the relationship between the percentage change in right striatal Glx and the pre-to-post change in connectivity strength for each identified cluster. Connectivity strength change was defined as post-stimulation minus pre-stimulation connectivity strength. Red and blue dots represent individual participants in the tTIS and sham groups, respectively. Solid and dotted lines indicate group-specific regression lines for the tTIS and sham groups, respectively.

**Table 2.** Clusters showing group differences in the association between changes in right striatal Glx levels and changes in seed-based functional connectivity.

| **Peak MNI**<br>**coordinates (x,y,z)** |  |  | **Cluster size,**<br>**voxels** | **Peak *t* value** | **Anatomical region** |
| --- | --- | --- | --- | --- | --- |
| 40 | -14 | -30 | 251 | 4.55 | Right fusiform gyrus |
| -34 | -8 | -42 | 236 | 5.04 | Left fusiform gyrus |
| 41 | -18 | 41 | 58 | 4.08 | Right postcentral gyrus |
| 28 | -26 | -22 | 53 | 3.93 | Right parahippocampal gyrus |
| 35 | -13 | 51 | 33 | 4.21 | Right precentral gyrus |

MNI, Montreal Neurological Institute. Clusters were identified using an uncorrected voxel-level threshold of $p < 0.0005$ combined with a cluster-extent threshold of $k > 30$ voxels. Peak $t$ values represent between-group differences in the regression slopes relating the percentage change in right striatal Glx levels to the pre-to-post change in seed-based functional connectivity.

## 4. Discussion

### 4.1. Modulation of striatal Glx following tTIS

The brain contains multiple large-scale circuits that coordinate neural activity across cortical and subcortical regions. Among these circuits, the cortico-basal ganglia-thalamo-cortical loop receives widespread cortical inputs and plays an important role in action selection, motor control, emotional regulation, and reward-based decision-making ^16, 43^. Within this loop, information processing is critically shaped by glutamatergic excitatory inputs and GABAergic inhibitory outputs. Glutamate, a major excitatory neurotransmitter in the brain, mediates excitatory inputs from the cortex and thalamus to the striatum, as well as thalamocortical feedback projections. In contrast, GABA, a major inhibitory neurotransmitter, is released by striatal projection neurons that project to the external and internal segments of the globus pallidus and to the substantia nigra, thereby contributing to the direct and indirect pathways of the basal ganglia. Through these pathways, the basal ganglia regulate inhibition and disinhibition within motor and associative circuits.

In the present study, right striatal Glx, a combined signal reflecting glutamate and glutamine, showed a significant pre-to-post decrease within the tTIS group. Because the present ^1^H-MRS protocol cannot separately quantify glutamate and glutamine, this finding should be interpreted as a change in the combined Glx signal rather than a glutamate-specific effect. Moreover, because the direct between-group comparison of Glx change did not reach statistical significance, this result should be considered suggestive rather than conclusive evidence of a stimulation-specific neurochemical effect.

The beat frequency used in the present study was 20 Hz, which falls within the beta frequency range. Beta-band activity has been associated with the maintenance of the current motor state and reduced flexibility in initiating new movements ^22, 44^. In this context, the observed decrease in striatal Glx may reflect changes in glutamate–glutamine metabolism associated with excitatory inputs to the striatum, rather than a direct reduction in synaptic glutamate release. However, because beta-band neural activity was not directly measured, it remains unclear whether the 20-Hz beat frequency modulated endogenous beta oscillations.

Recent large-scale electrophysiological studies have demonstrated that striatal activity reflects the topographical organization of cortical activity, indicating a close relationship between cortical inputs and striatal neural dynamics ^45^. In addition, a previous study reported cell-specific effects of tTIS, showing that excitatory pyramidal neurons are more sensitive to temporally modulated waveforms, whereas inhibitory interneurons show relatively preserved firing under pure high-frequency stimulation ^46^. Although these cell-specific effects were demonstrated in cortical regions and cannot be directly generalized to the striatum, they provide a potential mechanistic framework for interpreting the present striatal findings at the circuit level. Thus, the observed reduction in striatal Glx may reflect altered glutamatergic input-related activity or glutamate–glutamine metabolism within cortico-striatal networks.

tTIS has been proposed as a non-invasive approach to modulate deep brain structures, aiming to reduce direct engagement of overlying cortical regions. Although the absence of a significant between-group difference in Glx change precludes a definitive conclusion regarding stimulation-specific effects, the within-group decrease in Glx is compatible with the possibility that 20-Hz tTIS modulates glutamate-related metabolic processes in cortico-basal ganglia circuits. Nevertheless, this interpretation remains exploratory and requires confirmation in studies demonstrating a significant stimulation-by-time interaction and directly measuring the electrophysiological effects of stimulation.

### 4.2. Lack of detectable GABA+ modulation

No significant GABA+ changes were detected within either group or between groups. This finding may reflect the measurement characteristics of ^1^H-MRS rather than the absence of inhibitory circuit effects. ^1^H-MRS-derived GABA+ represents a broad tissue pool that includes co-edited macromolecular signals and does not directly index moment-to-moment synaptic inhibition ^47, 48^. Consequently, changes in GABAergic transmission, neuronal activity, or inhibitory synaptic efficacy may occur without a detectable change in bulk GABA+ levels. Previous studies have reported cortical GABA reductions after facilitatory non-invasive stimulation such as anodal tDCS ^49, 50^. However, the detectability of stimulation-induced GABA+ changes depends on stimulation parameters, acquisition timing, voxel location, tissue composition, spectral quality, and the sensitivity of GABA-edited ^1^H-MRS ^34, 49^. In addition, the magnitude and spatial distribution of tTIS-induced electric fields depend on the electrode montage and individual anatomy. Any striatal GABA+ change produced under the present conditions may therefore have been below the detection threshold.

A previous study showed that temporal interference stimulation can modulate hippocampal oscillations ^51^, but this does not imply that every beat frequency produces a measurable neurotransmitter change. Thus, the present null result should be interpreted as an absence of detectable bulk striatal GABA+ modulation, not as evidence that 20-Hz tTIS had no effect on inhibitory circuit function.

### 4.3. Association between striatal Glx changes and changes in resting-state functional connectivity

Within the tTIS group, greater reductions in right striatal Glx tended to accompany greater increases in functional connectivity between the right striatal seed and the sensorimotor cortex. Because the striatum receives dense glutamatergic input from sensorimotor regions and contributes to motor learning ^52, 53^, this pattern may reflect coupling between local neurochemical changes and cortico-striatal network dynamics. However, Glx does not directly measure synaptic glutamate transmission, and resting-state functional connectivity does not directly index synaptic plasticity. The findings therefore do not demonstrate long-term potentiation or establish the temporal or causal direction of the association.

Group-dependent Glx–connectivity associations were also observed in the right parahippocampal gyrus and bilateral fusiform gyri. The parahippocampal–hippocampal system contributes to episodic and contextual memory, and dynamic interactions between hippocampal and striatal systems have been reported during motor sequence learning and memory consolidation ^54–58^. The fusiform gyrus supports higher-order visual processing ^59^, and striatal connectivity with distributed cortical regions can vary during reward-related processing ^60^.

Together, these findings raise the possibility that the relationship between striatal Glx changes and functional connectivity extended beyond motor circuits to broader contextual, memory-related, and visual-associative networks. Nevertheless, the specific functional significance of the parahippocampal and fusiform findings remains speculative because no behavioral measures were obtained.

Overall, the results suggest group-dependent coupling between right striatal Glx changes and distributed connectivity changes. They should be interpreted cautiously because the analyses were correlational and used exploratory uncorrected thresholds, the direct between-group Glx difference was nonsignificant, and glutamate and glutamine could not be separated using the present ^1^H-MRS protocol.

### 4.4 Limitations and future directions

This study has several limitations. First, ^1^H-MRS measures bulk tissue signals and could not separate glutamate from glutamine; therefore, the observed Glx decrease cannot be interpreted as glutamate-specific or as direct evidence of altered excitatory synaptic transmission. Second, the spatial specificity of tTIS is limited, and contributions from overlying cortical regions cannot be excluded. Because the sham condition included 2,000-Hz carrier currents but no low-frequency interference envelope, the effects of temporal interference cannot be fully separated from those of the carrier currents without a no-stimulation control. Third, the absence of behavioral measures limits interpretation of the functional significance of the neurochemical and connectivity findings. Fourth, endogenous beta activity was not measured, and the pre–post design cannot determine whether Glx changes preceded connectivity changes or vice versa.

Repeated post-stimulation measurements, electrophysiological recordings, and multiple beat-frequency conditions are needed to clarify the temporal dynamics and frequency specificity of the effects. Fifth, the functional connectivity analysis used an uncorrected voxel-level threshold of p < 0.0005 with k > 30 voxels, increasing the likelihood of false-positive findings. Larger studies using more stringent multiple-comparison correction are required. Finally, dopamine and other neuromodulatory systems were not assessed, so their contributions to the observed changes remain unknown. Future studies should combine optimized or high-field MRS, individualized electric-field modeling, behavioral assessments, and both sham and no-stimulation control conditions to test the reproducibility and specificity of these findings.

## 5. Conclusion

A tTIS montage designed to target the striatum was associated with an exploratory within-group decrease in right striatal Glx and group-dependent associations between Glx and connectivity changes in sensorimotor, parahippocampal, and fusiform regions. These findings suggest potential coupling between neurochemical and network-level changes following 20-Hz tTIS but require confirmation in larger, well-controlled studies before clinical implications can be established.

## CRediT authorship contribution statement

**Shota Kimura**: Writing – original draft, Investigation, Formal analysis, Data curation, Visualization. **Hiroyuki Matsuta**: Writing – original draft, Formal analysis, Methodology, Data curation. **Kana Matsumura**: Data curation. **Fumiaki Iwane**: Formal analysis, Methodology. **Nobuhiro Hata**: Investigation, Resources. **Yoshiki Asayama**: Investigation, Resources. **Hisato Sugata**: Conceptualization, Methodology, Investigation, Formal analysis, Validation, Data curation, Writing – original draft, Writing – review & editing, Supervision, Project administration, Funding acquisition.

## Declaration of generative AI and AI-assisted technologies in the manuscript preparation process

The authors used AI for English-language editing. The authors reviewed and edited the content as needed and take full responsibility for the content of the article.

## Acknowledgements

The authors thank all participants who took part in this study.

